# Stitchr: stitching coding TCR nucleotide sequences from V/J/CDR3 information

**DOI:** 10.1101/2021.12.20.473544

**Authors:** James M. Heather, Matthew J. Spindler, Marta Herrero Alonso, Yifang Ivana Shui, David G. Millar, David S. Johnson, Mark Cobbold, Aaron N. Hata

## Abstract

The study and manipulation of T cell receptors (TCRs) is central to multiple fields across basic and translational immunology research. Produced by V(D)J recombination, TCRs are often only recorded in the literature and data repositories as a combination of their V and J gene symbols, plus their hypervariable CDR3 amino acid sequence. However, numerous applications require full-length coding nucleotide sequences. Here we present Stitchr, a software tool developed to specifically address this limitation. Given minimal V/J/CDR3 information, Stitchr produces complete coding sequences representing a fully spliced TCR cDNA. Due to its modular design, Stitchr can be used for TCR engineering using either published germline or novel/modified variable and constant region sequences. Sequences produced by Stitchr were validated by synthesizing and transducing TCR sequences into Jurkat cells, recapitulating the expected antigen specificity of the parental TCR. Using a companion script, Thimble, we demonstrate that Stitchr can process a million TCRs in under ten minutes using a standard desktop personal computer. By systemizing the production and modification of TCR sequences, we propose that Stitchr will increase the speed, repeatability, and reproducibility of TCR research. Stitchr is available on GitHub.

## Introduction

Alongside immunoglobulins, T cell receptors (TCRs) underly adaptive immunity in jawed vertebrates. They are the basis through which T cells initiate their major functions – detecting and responding to pathogens, cancers, and other threats – via recognition of peptides and other molecules displayed by MHC proteins. If congenitally absent, either from a lack of production of the proteins themselves or of the T cells that bear them, untreated individuals face high mortality risk during early infancy due to uncontrolled microbial infections(1, 2). Conversely, inappropriate recognition of molecules by TCRs can lead to autoimmunity or allergies(3). A lack of sufficient TCR-based responses to neoantigens and other tumor antigens contributes to the development and progression of malignancies, as illustrated by the clinical success of checkpoint blockade therapies in recent years(4, 5). It is hard to overstate the importance of T cell receptors within and beyond the field of immunology.

TCRs are produced through a process of somatic DNA recombination, in which gene segments are drawn from a panel at specific loci, the intervening DNA is imprecisely excised, and the once-separate coding portions are joined together(6). This process of V(D)J recombination – named after the variable (V), diversity (D), and joining (J) genes which can be rearranged together – is capable of producing an incredible diversity of TCRs. Combinatorial diversity is generated both by merit of TCRs being heterodimers of two polypeptides (pairing either alpha/beta or gamma/delta chains), with each polypeptide chain produced through this process, and by there being multiple V(D)J genes to be selected from at each TCR locus. Even greater diversity is introduced by the non-templated deletion and addition of nucleotides at the rearranged junctions. Ultimately, this process produces the region of the TCR gene that encodes the complementarity determining region 3 (CDR3), the hypervariable section of the TCR which contacts the antigen. Cumulatively, this system is estimated to be capable of producing ∼1 × 10^15^ unique alpha-beta TCRs in humans(7, 8), orders of magnitude greater than the number of T cells that an individual body could contain(9). Even though this ‘TCR space’ is not evenly utilized, there is still tremendous inter- and intra-individual receptor diversity, providing a substantial barrier to study. Investigation of the functional behavior of TCRs is additionally complicated by the diversity of their ligands. TCRs bind to short peptide fragments (derived from potentially any protein that finds its way into the body) presented in the groove of MHC proteins, which are among the most polymorphic genes in vertebrate genomes(10). Mechanistic studies on TCRs are therefore both extremely important, yet complicated due to the molecular diversity of the system.

TCR chains are frequently only reported as annotated rearrangements, consisting of the involved V and J genes plus the CDR3 sequence spanning the hypervariable rearrangement. In principle, assuming that the TCR has been correctly annotated, this contains all of the necessary information to reproduce the entire coding sequence (with the D gene sequence being contained within the CDR3 for beta chains). In order to perform experiments that require TCR expression, there is a need to reliably convert these concise TCR descriptions into full-length coding nucleotide sequences. To the best of our knowledge, no computational tool exists to do this. Instead, the traditional approach requires manual assembly of the V/J/CDR3 combinations of interest using germline TCR database repositories (such as IMGT-GENE/DB(11)) and a text editor or DNA software tool. While this approach can produce valid results, it is 1) slow and labor intensive, thus scaling poorly; 2) vulnerable to human error, leading to 3) poor repeatability (by one user) and reproducibility (by others)(12).

To overcome these limitations, we developed Stitchr, the first software capable of automatic generation of full-length coding human and mouse alpha-beta TCR nucleotide sequences from minimally reported V/J/CDR3 information. Stitchr produces a nucleotide sequence encoding the CDR3 inserted in-frame between the provided V and J genes segments, adding properly spliced upstream leader and downstream constant region sequences. The modularity of its approach also allows users to substitute in different, even non-natural, TCR gene segments, resulting in rapid sequence generation for protein expression and engineering experiments. We demonstrate that Stitchr produces the expected TCR sequences and verify the approach by synthesizing expression vectors for TCRs of known specificity and demonstrating their activity in Jurkat cells. We also report Thimble, a companion script which allows users to run Stitchr on single or paired-chain TCR repertoires, capable of processing a million TCRs in under ten minutes using a standard desktop personal computer. Finally, we illustrate case examples where high-throughput TCR datasets can be accurately converted to full-length coding equivalents. We propose that Stitchr and related tools will accelerate the pace of TCR research at the interface between experimental validation and high-throughput bioinformatic analysis.

## Materials and Methods

### Stitchr implementation

Stitchr was written in Python, tested exhaustively on Python 3.6.9 on Ubuntu and Python 3.7.7 on Mac OS. The only non-standard package needed for its operation is PySimpleGUI (>= version 4.45.0), if users elect to use the graphical user interface script. It is also provided with Thimble, a companion wrapper script which allows users to supply TCRs in a tab separated spreadsheet file, for the simultaneous stitching of multiple receptors. All scripts and data necessary to run Stitchr can be found on the GitHub repository: https://github.com/JamieHeather/stitchr.

By default Stitchr is equipped to generate human and mouse TCRs, using germline sequences downloaded from IMGT/GENE-DB(11) (last updated in 2021-08-22 and 2021-05-20 respectively). It takes as input a TCR rearrangement described by the V and J genes used, plus the CDR3 junction sequence, which can be provided as a nucleotide or amino acid sequence (running inclusively from the cysteine to the phenylalanine or equivalent residue) or as a longer nucleotide sequence extending further into the V/J genes.

The exact mode of determining the CDR3 junction depends on the input format. If an amino acid CDR3 is provided, Stitchr takes that sequence and looks for matching sequences with its N terminus and the C terminus of the translated relevant V gene, incrementally deleting V gene residues until it finds a match. The non-V portion of the CDR3 is then used to similarly search the translated J gene until the minimal CDR3 contribution is met, leaving the portion of the CDR3 junction which is not feasibly encoded by either germline gene (which will include any remaining residues from the D gene in the case of beta chains). The nucleotide sequences for the wholly-germline encodable V/J residues are then produced by trimming the provided sequences, and the intervening non-templated sequence is generated by selecting the most common codon for that residue from a species-specific frequency table. Users can also provide exact CDR3 junctions as nucleotides, which are first translated and then processed in a similar manner up until the non-templated region is determined, at which point the corresponding segment of the provided nucleotide sequence is used in place of a codon-optimized selection.

If users have additional nucleotide context beyond the junction, they can also use the ‘seamless’ stitching option for more faithful replication of the nucleotide sequence. In this mode the overlap detection occurs at the nucleotide level: the 5’ of the provided junctional sequence is searched against the incrementally deleted 3’ of the V gene, and the 3’ of the non-V portion is searched against the 5’ of the J. Once the sites of overlap between the V, the junction, and the J are determined, the extraneous sections of the germline V/J genes are removed and the sequences are joined. The leader and constant regions necessary for expression are then added. By default, these sequences will be inferred from the V and J genes chosen respectively, defaulting to the prototypical allele’s (*01) leader sequence if IMGT does not record one for the specified allele, and taking the relevant TRBC gene per TRBJ cluster (although both values can be optionally overridden). Users can also specify any additional sequences to the 5’ and 3’ of the total rearrangement (e.g. to add Kozak sequences, stop codons, restriction enzymes, or primer binding sites if TCRs are to be synthesized). If Thimble or the graphical user interface script is used, single stitched alpha and beta chains can be linked together with any desired sequence (e.g. a 2A self cleaving peptide sequence) for bicistronic vector expression.

### Benchmarking Thimble

Large TCR V/J/CDR3 datasets were obtained by randomly picking five samples from the Emerson *et al*. Adaptive Biotechnologies’ dataset (DOI: 10.21417/B7001Z, samples HIP02873, HIP02805, HIP02811, HIP02820, and HIP02855)(13) with the Python random.choice function, and downloading the entire VDJdb database (using the ‘vdjdb_slim’ file, accessed on 2021-02-02)(14). The Adaptive Biotechnologies data are beta chain sequences produced with a unified experimental and analytical pipeline from healthy donor peripheral blood mononuclear cell (PBMC) gDNA, while the VDJdb TCRs have been extracted and annotated from the literature and thus represent a diverse array of input cell sources and TCR identification processes, featuring both alpha and beta chains. Adaptive Biotechnologies uses custom non-standard identifiers to refer to TCR genes, so these were first converted to standard IMGT nomenclature and Adaptive Immune Receptor Repertoire Community (AIRR-C) format(15) with the Python script immunoseq2airr (version 1.2.0, DOI: 10.5281/zenodo.5224597). This was run with the following non-default parameters to account for input file formats and filter non-productive and non-interpretable rearrangements: -nd, -a, -or, -pf, -mf, and -p (pointing to the Emerson parameter conversion file provided in the repository). Rearrangements from both datasets were filtered to keep only in-frame potentially productive chains with both a V and J gene call. For ambiguous cases with >1 V gene call the first gene provided was used. Non-human TCRs were discarded from VDJdb. Note that Thimble successfully produces stitched sequences for >99% of all input V/J/CDR3 combinations, with the vast majority of those that fail lacking complete CDR3 junctions (i.e. they do not run from the conserved C to F residues, inclusively). In order to establish a wide dynamic range, variable numbers of TCRs were randomly drawn from these datasets (pooling all of the Adaptive files into a single resource) using the Python random.choices function, up- or down-sampling to 1e2, 1e3, 1e4, 1e5, or 1e6 rows, three times per dataset.

### Generating simulated TCR data with immuneSIM

50,000 AIRR-C compliant human TCR alpha and beta recombinations were simulated using the R package immuneSIM(16) (version 0.8.7), with a minimum CDR3 length of eight residues (and all other settings default). Rearrangements with CDR3 junctions not ending in one of the three conserved terminating residues found in human predicted-functional TRAJ/TRBJ genes (phenylalanine, tryptophan, or cysteine) were filtered and removed.

### High-quality long read TCR-seq dataset generation and analysis

In order to obtain ‘real’ full-length V domain TCR repertoire sequences, we leveraged published datasets produced in part by one of the authors previously, in which αβ TCR-seq was performed on RNA extracted from whole blood taken from 16 healthy volunteers(17, 18). These data were produced using a ligation-based 5’RACE strategy, in which random unique molecular identifier (UMI) barcodes were added to TCR cDNA prior to amplification, allowing for error-correction downstream. However, in contrast to previous studies, in which TCR rearrangements were annotated using only the constant region-proximal read of the paired-end sequencing (R1) prior to error correction, raw FASTQ were first merged using FLASH (version 1.2.11, default parameters)(19). This identifies paired end reads with 3’ overlap and combines them into a FASTQ with longer complete amplicons. Merged reads which share UMIs were then collapsed and error-corrected using stringent criteria: each UMI had to contain at least three reads, and the calls of those reads must all have quality scores >= Q25. The consensus base at each position was then determined, with the abundance of a given consensus reported by counting the number of associated barcodes, and outputed as FASTA files. While this greatly reduces the number of available reads in a repertoire file, the remaining reads are more likely to cover the entire variable domain and much less likely to contain PCR or sequencing errors. For the purposes of testing Stitchr/Thimble, these donor/locus specific FASTA files were combined into one large repertoire file.

Extended reads were then analyzed using a modified version of the TCR annotation software Decombinator(20, 21), called autoDCR. Like Decombinator, autoDCR uses short (20 nt) ‘tag’ sequences to populate an Aho-Corasick trie (or search tree) for efficient string matching based V/J gene identification. However, unlike the original Decombinator implementation in which single CDR3-proximal tags are selected for their unique occurrence in single genes, autoDCR tiles 20-mer tags overlapping 10 nt across the entirety of every allele of every gene, making V and J gene calls based on the presence of multiple tag matches. While the much larger trie takes longer to search each read, it outputs V and J gene calls with allele-level accuracy. This allows determination of sequence across the length of the rearrangement, enabling selection of reads which include the start of the V gene. Technically, this is achieved by using the ‘-jv’ flag, which outputs the ‘jump’ values indicating the furthest positions of V/J tag matches: filtering on v_jump values = 0 selects reads where the first tag match corresponds to the start of the V-REGION. Additionally, TCRs with ambiguous gene calls (>1 allele), or those using alleles for which only partial nucleotide information is available in IMGT, were filtered out.

This feature of detecting overlapping tags was also utilized to perform rudimentary detection of novel TCR alleles, guided by the hypothesis that a) individuals with alleles not present in IMGT will exist in our cohort and b) most of these are likely just single nucleotide variants (SNV). TCRs with fully sequenced variable domains from each donor were screened for potential novel alleles, indicated by multiple rearrangements using the same V gene but which all share a two-tag mismatch with the reference (as a SNV relative to a recorded germline gene will result in two consecutive overlapping tags failing to match), with the same sequence spanning the break. To distinguish potential novel allele variants from PCR/sequencing errors, sequences had to meet the following criteria: 1) be present in a V gene with >= ten distinct recombinations with unambiguous gene calls; 2) account for >= 5% of the reads belonging to that V gene; 3) be found in >= three unique recombinations; 4) account for >= 10% of reads for that V gene which contain a break of two tag matches. As an additional check, we called potential novel alleles only if they occurred in the top two most abundant sequences for that gene, which would then constitute the genotype for that gene in that donor. Inferred potential novel alleles were assigned an identifier indicating their variant suffixed to their original reference allele match (e.g. TRAV27*01_A233G) and output as FASTA reads in IMGT format, and then either appended to the IMGT database to re-generate tags for autoDCR, or supplied to Stitchr by including them in the ‘additional-genes.fasta’ file.

### In vitro TCR validation

Full-length TCRα and TCRβ coding regions generated by Stichr were used to generate TCRα-TCRβ lentiviral expression constructs with the BioXp 3200 system (Codex DNA) as previously described(22). In brief, the TCRα and TCRβ sequences were generated with Stitchr (using the same TRBC2 constant region and TRAV21/TRBV6-5 leader sequences for reliable expression) and cloned into a pReceiver-based lentiviral vector (GeneCopoeia) that contained an EF1α promoter and puromycin resistance gene. The bicistronic TCRα-TCRβ coding region incorporated a P2A ribosomal skip motif(23) to generate independent TCR polypeptide chains. Lentivirus was packaged into VSV-G pseudotyped particles using the third-generation ViraSafe Lentiviral Packaging System (Cell Biolabs) and Lenti-Pac 293Ta cells (GeneCopoeia)(24). Fresh lentiviral supernatant was used to transduce TCRβ-deficient (ΔTCRβ) Jurkat cells (J.RT3-T3.5; ATCC TIB-153), which were previously engineered to stably express human CD8 (lentiviral construct layout: PGK promoter - CD8A-P2A-CD8B(M-1) – IRES-blasticidin resistance gene)(25, 26). Transduced ΔTCRβ Jurkat cells were selected with puromycin for 14 days and introduced TCR surface expression confirmed by antibody staining for CD3 and TCRαβ surface expression.

TCR-engineered Jurkat cells and target cell lines (**Supplementary Tables 1 and 2**) were cultured in RPMI 1640 supplemented with 10% FBS, 2 mM glutamine, and penicillin/streptomycin. Target cell line HLA type information was obtained from the TRON Cell Line Portal(27). PBS-washed target cells were stained with 1 μM CFSE (eBioscience) for 10 min at RT, before being quenched with 5X volumes of complete media, incubated on ice for 10 min, and repeatedly washed in media. Stained cells were then peptide pulsed with 10, 1, or 0 μg/ml of the relevant peptide epitope (GenScript, >= 85% purity) for 1 hour at RT in complete media, washed twice, and co-cultured overnight (18-22 h) with Jurkat cells expressing the relevant TCR at an effector-to-target ratio of 2:1. Experiments were set up in round-bottomed 96 well plates, with 2e5 Jurkats and/or 1e5 target cells per well, with each experimental condition in triplicate. Plates contained additional no target cell negative control and anti-CD3 antibody (CD3-2, Mabtech) positive control wells. Following co-incubation, cells were washed in PBS and then FACS buffer (PBS with 2% FBS and 1 mM EDTA). Cells were then stained in the dark on ice for 30 min with 0.5 μl anti-CD62L and anti-CD69 antibodies conjugated to PE and APC respectively (BioLegend, clones DREG-56 and FN50) in 100 μl FACS buffer per well. After washing twice, cells were resuspended in 100 μl of FACS buffer with 2 μl 7AAD (BioLegend) and incubated in the dark on ice for a further 5 minutes. Antibody staining was quantified on an Accuri C6 Plus flow cytometer (BD Biosciences) with a CSampler. FCS data were analyzed with FlowJo version 10.7.1.

### Data analysis and visualization

All non-FCS analysis and data visualization was carried out in Python >= 3.6.9, using a combination of matplotlib (version 3.3.2)(28), pandas (1.1.2)(29), and seaborn (0.11.0)(30). PDB TCR amino acid sequences were aligned using Clustal Omega(31) version 1.2.4, accessed via the web portal in April 2021.

## Results

We designed Stitchr to take the minimal V gene, J gene, and CDR3 junction sequences typically used to report TCR chains and generate a complete functional coding nucleotide sequence (**Figure 1A**). Stitchr accommodates multiple CDR3 formats: the junction sequence (running inclusively between the conserved V gene cysteine and J gene phenylalanine) can be supplied in either amino acid or nucleotide form. Stitchr determines which residues of the translated CDR3 can be encoded wholly by germline sequences, and then all 5non-templated5 nucleotide sequences between those portions (which also contains D gene contributions for the beta chain) will either be generated from a species-specific codon frequency table (for amino acid input - **Figure 1B**) or will be cropped out of the provided nucleotide sequence (for nucleotide input - **Figure 1C**). Alternatively, if a nucleotide sequence that extends beyond the edges of the CDR3 junction is supplied, Stitchr seamlessly integrates into the germline V and J genes after computationally ‘deleting’ from the germline genes and looking for overlapping sequences (**Figure 1D**). Complete TCR nucleotide sequences (**Figure 1E**) are output after splicing on 5’ leader and 3’ constant region sequences, with the option to add arbitrary sequences to either side of a gene (e.g. for gene regulatory or PCR/cloning purposes). Stitchr can be run as a command line tool for a single chain recombination, or via a wrapper script (discussed below) for larger numbers of sequences, which also supports the production of bicistronic sequences for paired alpha and beta chains (e.g. for making expression vectors). Alternatively, we designed a graphical user interface that can be used to generate single or paired chain receptors (**Supplementary Figure 1**).

**Figure 1.**
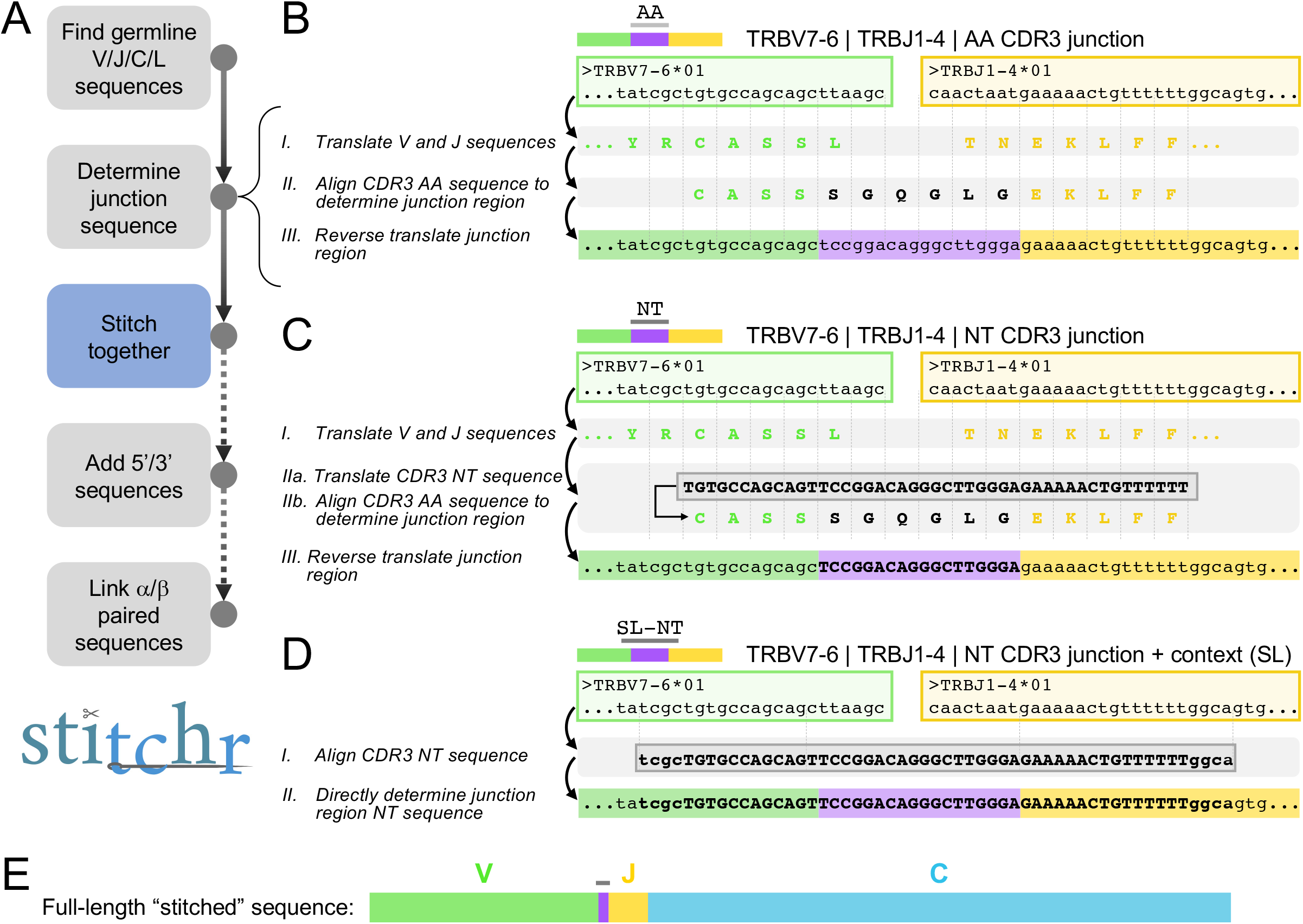
Schematic of Stitchr algorithm. **A**, Overview of Stitchr modules. Stitchr first obtains germline V gene, J gene, constant region (C) and leader (L) sequences from IMGT/GENE-DB. Next, the junction-spanning sequence is determined, depending on input mode (see **B**-**D**), and the complete TCR sequence is assembled. Complete single chain rearrangements can subsequently have arbitrary user-provided sequences appended to the 5’ or 3’ of the TCR, and finally alpha/beta chain pairs can be combined (e.g. via a 2A self-cleaving peptide sequence) into a bicistronic single expression sequence. **B**, When an amino acid (AA) CDR3 junction sequence is provided, the V and J genes are translated (I), aligned, and ‘deleted’ back from the CDR3-proximal edge until the longest possible overlap with the appropriate side of the junction is found (II), i.e. the longest suffix of the V that matches the prefix of the CDR3, or vice versa for the J. The remaining residues which cannot be encoded by the germline genes are then ‘reverse translated’ using a codon frequency table (III), and the trimmed germline genes and non-templated residues are concatenated. Vertical dotted lines show codons. **C**, If provided a nucleotide (NT) CDR3 junction sequence (depicted by bold/capitalized font), the germline genes are again translated (I), as well as the CDR3 sequence (IIa). The amino acid sequences are aligned and the germline contributions to the CDR3 are determined (IIb). The AA sequence is then converted to NT, however instead of assigning codons for the non-templated residues based on a codon usage table, the nucleotides in the provided CDR3 are used (III, bold text indicates retained original NT sequence). **D**, If the provided junction sequence includes additional nucleotide sequence context that extends beyond the CDR3 (depicted by lowercase text), the ‘seamless’ (SL) option can be used. In this mode, V and J germline genes are again deleted to the edge of the overlapping NT sequence (vertical dotted lines), allowing Stitchr to seamlessly combine germline V and J with the provided CDR3-spanning sequence (II). All three options produce a full-length TCR sequence (**E**) that encodes the same amino acid sequence, with the seamless option reproducing the identical nucleotide sequence (assuming the correct V and J alleles were provided).

To test the capabilities of Stitchr, we used the command line interface to generate full-length TCR sequences from four published receptors that have rigorous data demonstrating epitope recognition, including solved structures of the TCR-peptide-MHC complex, and which cover a range of V/J gene combinations and HLA restrictions (**Supplementary Tables 1 and 2**)(32–35). We then downloaded amino acid sequences for each of these alpha-beta TCRs from the Protein Data Bank (PDB)(36) and aligned them against translations of the Stitchr-generated sequences (**Figure 2A**). With one exception, the variable domain sequences produced by Stitchr aligned perfectly with the PDB TCR structures. The exception – in the alpha chain V gene segment of the MAG-IC3 TCR – is explained by that TCR having been engineered to include a non-germline modification in order to increase affinity(35). As Stitchr makes use of a modular approach in its selection of genes, alternative and additional sequences can be added to expand the types of TCRs that can be produced. When we supplied Stitchr with a suitable reference for this altered V gene, it accurately reproduced the PDB amino acid sequence. Thus, Stitchr faithfully replicates TCR sequences, at least when the templated sections are present in the ‘germline’ genes supplied to it. Stitchr’s potential for rational protein design was further explored by generating the beta chain of the anti-MART1 TCR DMF5(37) in combination with different constant regions. Stitchr generated appropriate coding sequences correctly spliced onto human TRBC1, TRAC, TRDC, and TRGC1, and mouse TRBC1 constant regions (**Supplementary Figure 2**), thus supporting the use of Stitchr for generating even non-natural TCR sequences.

**Figure 2.**
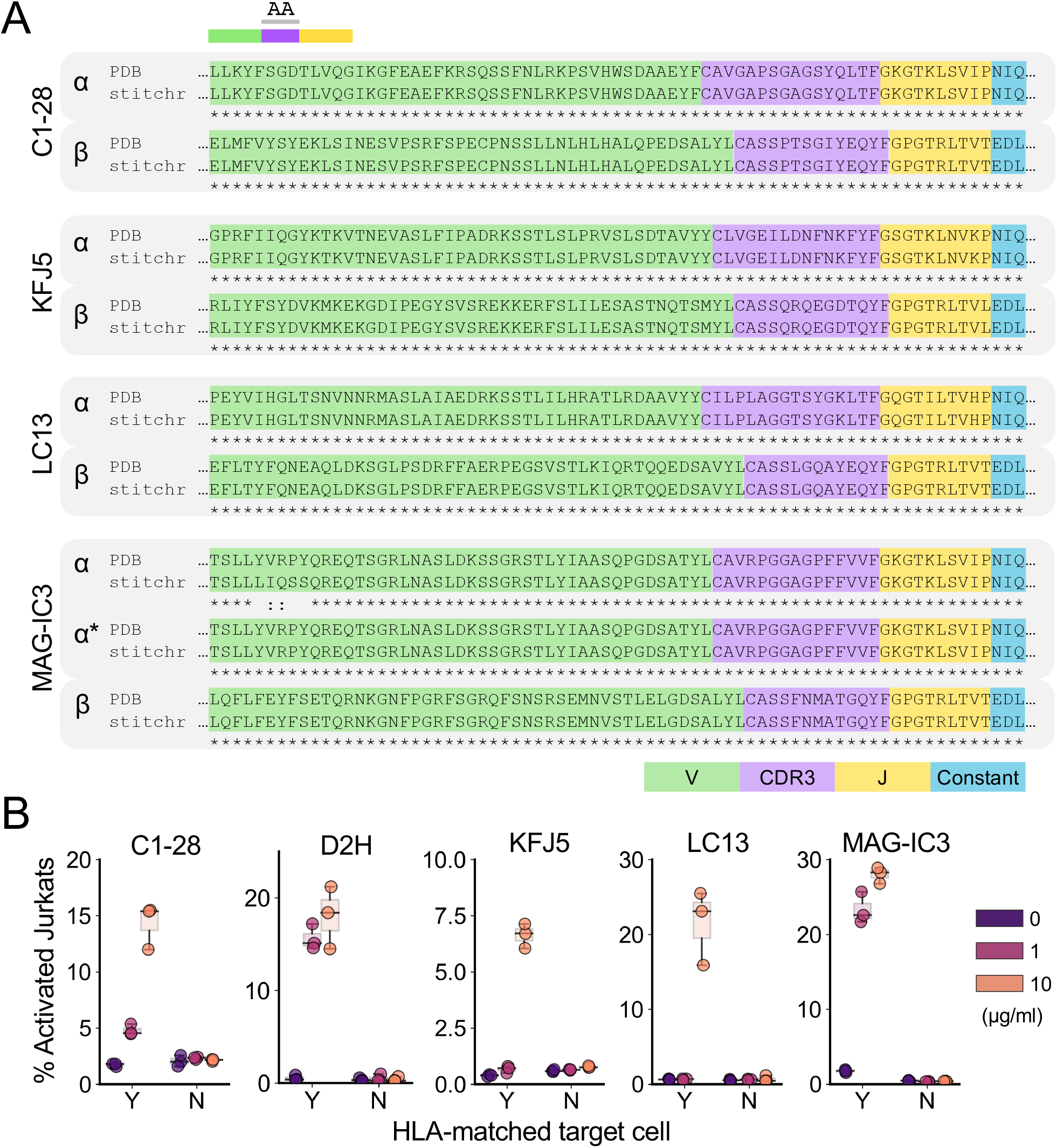
Validation of Stitchr-generated TCR sequences. **A**, Amino acid sequences of four TCR heterodimers were extracted from PDB structures and aligned against Stitchr-produced sequences for the same rearrangements (using ‘ATG’ in place of leaders omitted from the crystallized structures), showing the correct incorporation of junction sequence and constant region. MAG-IC3 ‘α*’ sequence indicates Stitchr output using a modified TRAV21*02 gene to replicate the engineered amino acid sequence used in the PDB structure. **B**, Functional validation of Stitchr-produced TCR sequences using a Jurkat activation assay. CD8-positive, TCRb-negative Jurkat cells were transduced with one of five different TCRs and co-cultured with peptide pulsed (10, 1, or 0 μg/ml) HLA-matched or mis-matched target lines. Data shown are triplicate technical replicates from one experiment and are representative of at least two independent biological repeats.

To test whether TCRs produced by Stitchr are functional, we generated expression constructs for the four TCRs from **Figure 2A**, plus an additional published TCR which expanded the range of TCR genes and HLA alleles covered (but which lacked structural data)(38) (**Supplementary Tables 1 and 2**). P2A-linked bicistronic TCR constructs were stably expressed in ΔTCRβ Jurkat cells and co-cultured with cognate peptide-pulsed cancer cell lines that express the relevant HLA allele. Jurkat activation (CD69+/CD62L-negative) was assayed by flow cytometry (**Supplementary Figure 3**). We observed dose-dependent peptide-induced activation of TCR-Jurkat cells when cultured with target cells bearing the appropriate HLA allele, but not with HLA-mismatched target cells (**Figure 2B**). These results confirm that Stitchr generates functional TCRs that reproduce the antigen specificity of the rearrangements they replicate.

**Figure 3.**
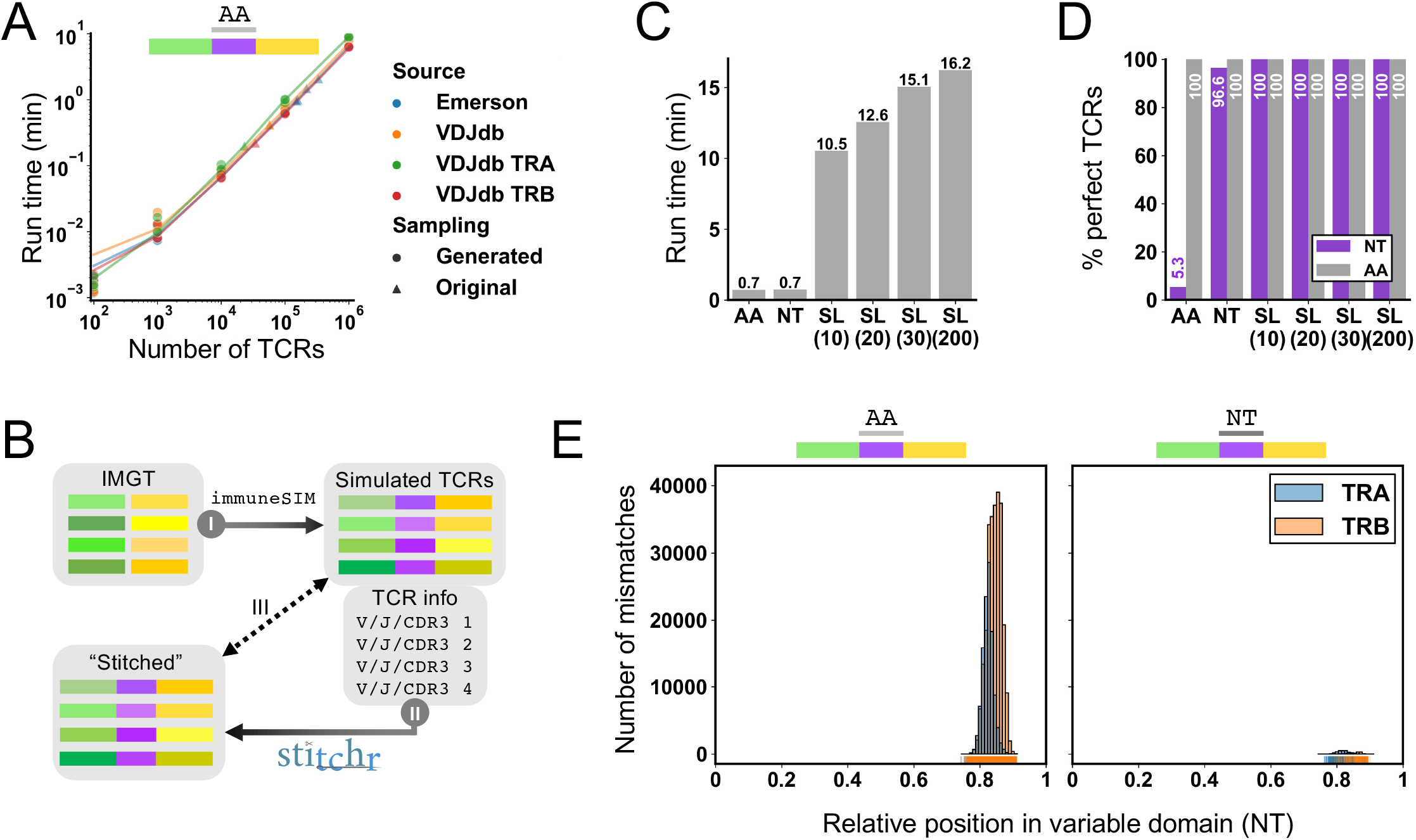
Application of Stitchr to high-throughput TCR datasets using the companion script Thimble. **A**, To benchmark the speed of Thimble, large TCR datasets with amino acid CDR3s provided were downloaded either from bulk beta chain TCR-seq datasets^13^, or from the curated antigen-associated TCR database VDJdb^14^ (processed both all together and by each chain individually). Thimble, the high-throughput interface to Stitchr, was run on these original files (triangle markers), and from files containing 100-1,000,000 TCRs generated by randomly re-sampling these files (dot markers), with each repertoire size randomly produced 3 times. Connecting lines indicate bootstrapped locally weighted linear regressions. **B**, Overview of sequence-level Stitchr validation. TCRs with known V/J/CDR3 information and nucleotide sequence were produced by *in silico* recombination of IMGT-stored germline genes using immuneSIM (I). V/J genes and CDR3 information (taken as exact junctions in nucleotide or amino acid forms, or as nucleotides with additional padding sequences for seamless mode) were input to Stitchr (via Thimble) (II). TCR variable domain sequences produced by Stitchr were then compared against the corresponding parental simulated TCR sequences (III). **C**, Run time duration of Thimble applied to 50,000 α and β TCRs generated by immuneSIM, comparing different formats of junction region input: amino acid (AA), nucleotide (NT), nucleotide with padding nucleotides 5’ and 3’ for seamless (SL) integration, either 10, 20, 30, or 200 (200 5’, 30 3’). **D**, Percentage of TCRs produced by Stitchr for which the variable region (start of V gene to end of J gene) perfectly matched the input sequence generated by immuneSIM, at both the nucleotide (NT, purple) and translated (AA, grey) levels. **E**, Histogram of positional mismatches between simulated and stitched sequences for NT and AA junction input modes. Histograms were generated with 111 bins, so each bar corresponds approximately to one codon (given the variable domain length distribution of ∼333 nucleotides, **Supplementary Figure 5A**).

With the rise of high-throughput TCR sequencing technologies, large TCR datasets are increasingly available. However, many such studies do not sequence the entire variable domain, and an even smaller fraction sequence the entire transcript. Even when the entire chain is sequenced, it is not often reported nor sufficient raw data provided to extract it, which can limit usefulness for certain applications. To remedy this limitation, we developed Thimble, a companion wrapper script which allows Stitchr to be applied to multiple TCRs in a single command. We benchmarked Thimble using large, published datasets (see Methods). This revealed that run-time scales linearly with number of input TCRs, taking under ten minutes to process a million TCRs on a standard desktop personal computer when provided with amino acid CDR3s (**Figure 3A**), successfully stitching >= 99.96% of all input rearrangements (**Supplementary Figure 4A**). The Emerson *et al*. data(13) also had CDR3-proximal read data available, and when used as input for seamless mode stitching, resulted in ∼10x times slower run-time relative to the amino acid sequence input (**Supplementary Figure 4B**).

**Figure 4.**
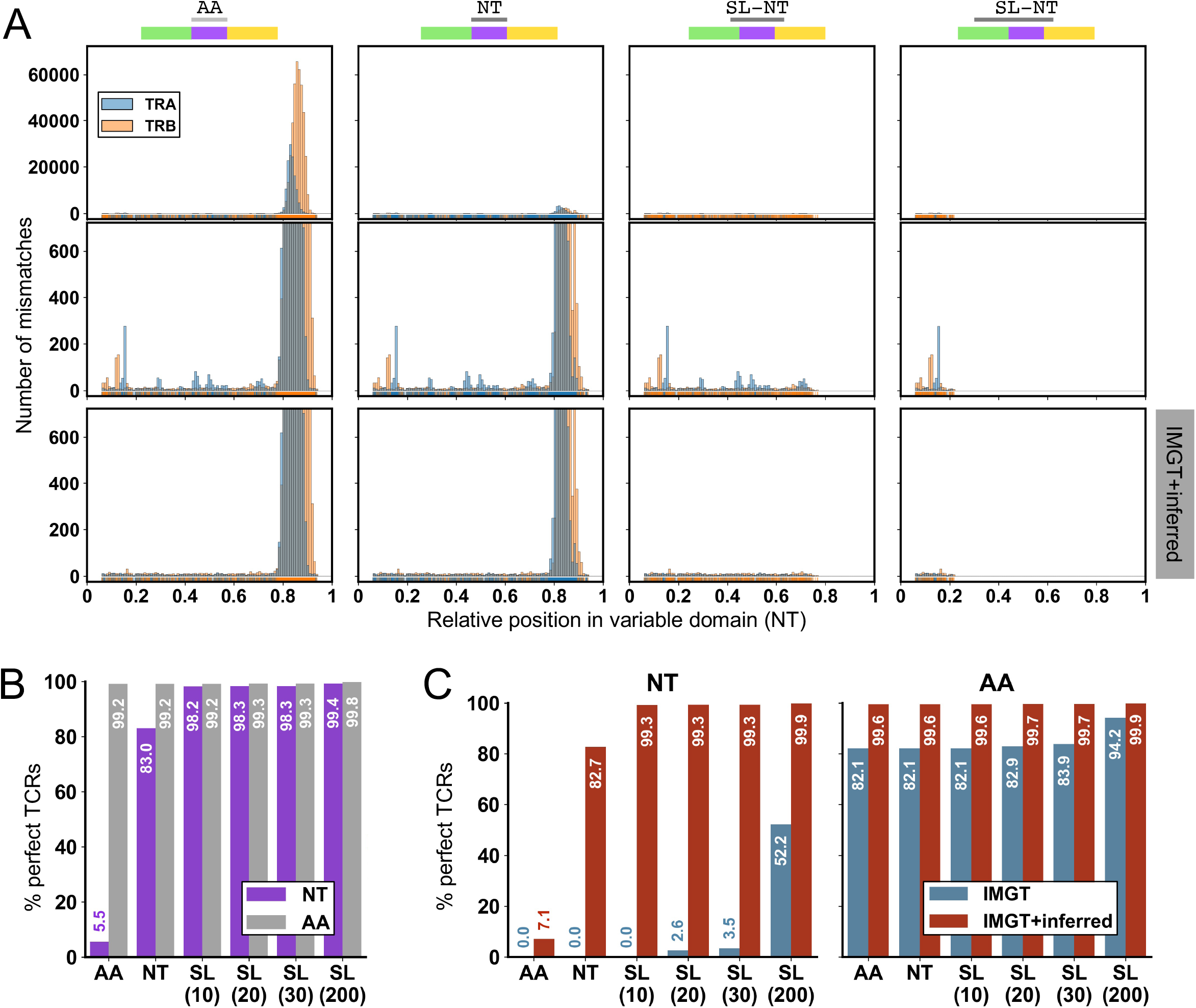
Assessment of Stitchr/Thimble accuracy on high-throughput TCR-seq data. **A**, Relative positional mismatches between Thimble-generated sequences and original input sequenced TCRs for different junction inputs: (columns left-to-right): AA, NT, SL (20), SL (200). Top row shows errors when using IMGT-provided TCR germline genes only. Middle row shows the same analysis with expanded Y-axis to highlight the bottom hundredth of the mismatch range. Bottom row shows mismatches upon rerunning Stitchr/Thimble when providing additional novel TCR alleles inferred from the individual donor repertoires. **B**, Percentage of all TCRs produced by Stitchr/Thimble that match perfectly to the original input sequences, using the IMGT reference. **C**, Percentage of only those TCRs that use a potentially novel inferred V gene allele and agree perfectly between TCR-seq and Stitchr/Thimble output, before (blue) and after (red) including those alleles in the reference dataset, at the nucleotide (left) or amino acid (right) level.

While these published datasets allowed us to benchmark basic run results, they do not contain complete variable domains, and were thus unable to confirm that Stitchr and Thimble were generating the correct nucleotide sequences. To generate gold-standard TCR sequences spanning the entirety of the variable domain necessary to rigorously assess this, we used the immuneSIM tool(16) to simulate V(D)J recombination, creating known TCR sequences from the IMGT germline reference database and predetermined generation probabilities (**Figure 3B**). This produces a repertoire with a normal distribution of variable domain lengths (**Supplementary Figure 5A**). These were then converted to Thimble input files, submitting the CDR3 junction as either amino acids (AA), nucleotides (NT), or as nucleotides with different 5’ and 3’ lengths (10/10, 20/20, 30/30, or 200/30 nt) for seamless (SL) integration. As expected, the seamless options took longer to run, scaling with longer nucleotide contexts (**Figure 3C**), but all options were completely ‘stitchable’ (**Supplementary Figure 5B**). By comparing the nucleotide and translated sequences produced by Stitchr to the original sequences generated by immuneSIM we observed that Stitchr’s accuracy was very high, perfectly recapitulating AA sequences in all modes, NT sequences in all seamless modes, and almost 99% correct NT sequences even when supplying CDR3s as AA input (**Figure 3D and Supplementary Figure 5C**). Examination of the distribution of these mismatches between input and output sequences revealed that they were all confined around the relative positions 0.8-0.9, corresponding to the expected location of the CDR3 where Stitchr generates codon-optimized sequences to fill in non-templated regions (**Figure 3E**). Note that providing the junction as a NT sequence still infrequently produces differences, as sometimes V(D)J recombination will delete only part of the codon(s) at the recombining edges before adding alternate nucleotides that still encode the same amino acid, while Stitchr defaults to using the germline-encoded sequence (e.g. **Figure 1C**, where the last residue of the ‘CASS’ motif was encoded by ‘AGT’ in the rearrangement but the ‘AGC’ found in the germline gene gets used).

Synthetic sequences do not necessarily reflect the true complexity of empirically sequenced repertoires. We therefore leveraged a published TCR-seq data generated with a unique molecular index (UMI)-barcoded 5’ RACE protocol, allowing production and allele-level annotation of stringently error-corrected TCR rearrangements running from the start of the V gene region to the end of the J (see Methods). This produced ∼365,000 TCRs of known sequence that were then submitted to Stitchr/Thimble, providing the CDR3 junction in different formats as with immuneSIM (**Supplementary Figure 6A**). Basic Stitchr results were broadly similar to those seen for the simulated data (**Supplementary Figure 5D-F**). Inspecting the accuracy profiles revealed that as expected, the majority of cases produced correct amino acid sequences, with some expected nucleotide mismatches when providing CDR3 junctions as AA or NT (**Figure 4A-B**). However, there were some additional nucleotide mismatches even when providing extended junctions for seamless integration **Figure 4A**, top two rows) that occurred outside of the region included in the padded junction (which gets integrated into the stitched sequence and is thus perfectly matched). While some of these mismatches are likely technical errors accumulated during the TCR-seq protocol (during RT, PCR, or sequencing) that survived the error-correction process, some appeared at markedly higher frequencies than others (**Figure 4A**, middle row). We hypothesized that some of these peaks may represent novel TCR alleles present in our cohort that aren’t represented in the IMGT reference database. We performed a novel allele inference analysis on each of our donor repertoires (see Methods and **Supplementary Figure 6B**) and introduced the potential novel alleles inferred from that process both to our TCR annotation software and Stitchr reference files. When we re-ran the analysis, the largest mismatch peaks at both the NT (**Figure 4A**, bottom row) and AA (**Supplementary Figure 5G**) levels were no longer present. When we restricted the analysis to TCRs that were found to incorporate a potentially novel allele, we observed an increase to near-completely perfect amino acid TCR replication when using the combined IMGT+inferred allele reference database (**Figure 4C**). Collectively, these results demonstrate that Stitchr can be scaled to accurately generate full-length TCR sequences for high-throughput datasets, using a variety of input CDR3 formats.

## Discussion

TCRs have been intensely studied since their discovery in the 1980s, drawing on many innovative approaches to overcome the challenges presented by their complexity. In particular, the advent of high-throughput sequencing technologies (TCR-seq) has allowed the identification of many orders of magnitude more rearranged TCRs than were possible with traditional techniques(39). More recently, microfluidic and other single-cell technologies have enabled high-throughput pairing of alpha-beta chain information through sequencing(40, 41), and even high-throughput functional cloning of screenable libraries(22). From these efforts, and those of other experimental approaches, there now exist various databases of deep-sequenced TCR repertoires(42–45), antigen- or pathology-associated TCRs(14, 46, 47), and structurally determined TCR-pMHC interactions(48–50). These provide a wealth of information for other researchers to build upon. Moreover, the translational potential of TCRs is increasingly being explored, particularly for anti-cancer treatments, as TCRs are capable of directly targeting proteins other than those expressed on the cell membrane (in contrast to monoclonal antibodies, for example). This includes a range of both cellular (TCR-T) and soluble (e.g. ImmTACs and other TCR fusions) TCR therapies undergoing clinical trials(51–53). The ability of TCR research to effect change in basic immunology and in the clinic has never been greater.

Despite these advances, barriers remain both within and between the sub-fields of TCR biology. Many of these relate in some way to one of two problems: 1) researchers are often not working with full-length TCR sequences, and 2) the methodologies of different fields tend to work at different scales. The former issue typically arises as researchers work off sequencing reads shorter in length than TCR variable domains, or from reported TCRs in which the sequence has been converted to the detected or inferred V and J genes and CDR3 sequence used(54). Both issues can present a barrier to many subsequent experimental and computational applications, such as synthesizing and expressing TCRs for validation of antigen specificity, or computationally investigating the contribution of different regions of the TCR to certain biological properties. TCRs proposed to enter pre-clinical testing require empirical validation of their specificities, due to effects like bystander activation(55), non-specific MHC multimer reagent staining(56, 57), and cross-reactivity(58, 59). Moreover several bioinformatic strategies to predict antigen specificity from sequence make use of information encoded at regions outside those typically sequenced in CDR3-centric protocols(60–62).

Here we introduce Stitchr, a python script that uses the V/J/CDR3 TCR information commonly used to report TCR identity and a table of germline sequences to generate corresponding full-length coding nucleotide sequences. We show that the sequences produced by Stitchr faithfully reproduce the amino acid sequences of the TCRs they aim to replicate. Stitchr can also be used to assemble TCRs with non-natural sequences, as may be desirable in TCR engineering applications. Moreover, through use of the companion script Thimble, we demonstrate that Stitchr is capable of processing a million sequences in ten minutes on a standard desktop personal computer, meaning that deep-sequencing data covering the CDR3 portion of TCRs can be converted into full-length sequences in a high-throughput manner.

The modular approach by which Stitchr reads in and assembles TCR sequences provides an effective way to generate edited or non-natural TCR sequences. A common modification is the use of alternative constant regions, typically to reduce the likelihood of mispairing with endogenous TCRs when introducing an exogenous TCR. This can take the form of swapping either whole(63, 64) or partial(65, 66) constant regions with their orthologous equivalents from different species (e.g. swapping human for mouse sequences), swapping whole or partial sequences between loci within a species (e.g. swapping whole(67) or partial(68) alpha/beta constant region sequences), or swapping alpha/beta chain regions for gamma/delta TCR equivalents(69)). Constant region domain swaps or supplementation of modified variable region genes are simply performed in Stitchr by including the wanted sequence in the reference data (**Supplementary Figure 2** and **Figure 2A** respectively). Arbitrary non-TCR sequences can also be appended to either end of a TCR rearrangement, or even used to bridge an alpha and a beta into a bicistronic sequence, to facilitate molecular manipulations and transgene expression (as used for the TCRs tested in **Figure 2B**). It is also increasingly appreciated that the IMGT database of germline TCR alleles is incomplete – for example, a recent pre-print reports discovery of 38 novel TRBV alleles mined from public datasets(70) – which raises concerns about applicability given the over-representation of certain geographic populations in these public datasets(71). Much as with non-natural modifications, we show that Stitchr can be used to include novel inferred alleles in the sequences it produces, improving the fidelity of the TCRs it generates (**Figure 4**).

One potential limitation of Stitchr is that it can only produce TCRs using the available component sequences (V and J genes plus leader and constant regions). While additional sequences can easily be provided, it is also possible that the V/J/CDR3 information being used as input may not faithfully reflect the ‘true’ sequenced TCR, e.g. if the TCR contained polymorphisms which were not captured by CDR3-proximal sequencing. V gene polymorphisms can impact upon antigen recognition(72) and surface expression levels(73), and could theoretically be recognized as foreign antigens themselves(58), thus such differences could prove functionally relevant depending on the intended application. Therefore, we recommend that wherever faithful reproduction of TCR sequences is required, full-length variable domain sequencing is performed. Advanced users may draw on their own sequencing data, along with databases of inferred TCR alleles like OGRDB(74) and VDJbase(75) to supplement or replace the provided germline sequences as needed.

While we have demonstrated the construction and validation of a small number of TCR expression constructs here, we anticipate that Stitchr and Thimble will prove useful in large-scale TCR gene synthesis and validation efforts. The domain switching illustrated in **Supplementary Figure 2** could be adapted and expanded to any number of TCR-related efforts, e.g. to convert TCRs to soluble forms by adding appropriate constant region sequences(76). As screens for cancer or infection-recognizing clonotypes increase(77), and more engineered TCR assays and immunotherapies are developed(51, 52), we believe that a tool like Stitchr stands to benefit the field by reducing the time and effort currently spent manually assembling full-length TCR expression construct sequences. Moreover, by effectively converting TCR design into a programmatic process it becomes exquisitely repeatable and reproducible, consistently producing the same output given the same input. This will contribute to rigor in TCR research, accelerating the pace and minimizing the chances of mistakes in TCR sequence production, both in basic research and potentially in the clinic.

## Data and code availability

Healthy donor UMI-barcoded merged FASTQ files are available on SRA under the accession PRJNA359580. Raw flow cytometry data from Jurkat validation experiments are available from FlowRepository, under the experiment IDs #4949-4958 inclusively.

Stitchr is available on GitHub under a BSD 3-Clause License here: https://github.com/JamieHeather/stitchr. The immunoseq2airr script for converting Adaptive Biotechnologies data to standard IMGT nomenclature and AIRR-C format (https://github.com/JamieHeather/immunoseq2airr) and the autoDCR script for TCR annotation (https://github.com/JamieHeather/autoDCR/) are similarly available. The code underlying the analyses shown in this manuscript (plus additional input data) is available here: https://github.com/JamieHeather/stitchr-paper-analysis.

## Acknowledgments

JMH would like to thank David Pinto (under the pseudonym Carnë Draug) and Margaret Axelrod for their early feedback on and contributions to the code on GitHub, and both Margaret and Justin Balko for their encouragement and feedback in adapting Stitchr for murine TCRs.

## Author Contributions

JMH: Conceived/wrote software, identified/verified TCRs, drafted manuscript

MHA: Verified TCRs

MJS: Identified/verified TCRs

YIS, DGM: Guided experimental strategy, performed preliminary experiments

DJ, MJS, MC, ANH: Project oversight, supervision, and sourcing funding

All authors: Contributed to and critically appraised experimental design and manuscript

