## Supplementary Tables and Figures for "Stitchr: stitching coding TCR nucleotide sequences from V/J/CDR3 information"

| TCR clone | HLA allele | Peptide | Source | HLA-matched target line | Publication DOI | PDB accession |
|---|---|---|---|---|---|---|
| **MAG-IC3** | A*01:01 | EVDPIGHLY | MAGE-A3 (Human) | MDA-MB-436 | 10.1038/srep18851 | 5BRZ |
| **C1-28** | A*24:02 | RFPLTFGWCF | Nef (HIV) | SW620 | 10.1038/srep03097 | 3VXM |
| **KFJ5** | B*07:02 | APRGPHGGAASGL | NY-ESO-1 (Human) | Karpas-299 | 10.1038/s41467-018-03321-w | 6AVF |
| **LC13** | B*08:01 | FLRGRAYGL | EBNA2A (EBV) | MDA-MB-436 | 10.1016/S1074-7613(02)00513-7 | 1MI5 |
| **D2H** | A*11:01 | GTSGSPIVNR | NS3 (DENV) | Karpas-299 | 10.1038/ni.3850 | - |

**Supplementary Table 1**: Antigen and reference details for the TCRs used for low-throughput Stitchr validation in Figure 2. Cell lines that do not express the relevant HLA class I allele were used as the HLA negative controls for other TCRs, as well as additional cell lines (e.g. SK-N-SH and SU-DHL-1, which were used as negative control targets for the KFJ5 TCR).

| TCR clone | TRAV | TRAJ | TRA CDR3 | TRBV | TRBJ | TRB CDR3 |
|---|---|---|---|---|---|---|
| **MAG-IC3** | TRAV21*02 | TRAJ28 | CAVRPGGAGPFFVVF | TRBV5-1 | TRBJ2-7 | CASSFNMATGQYF |
| **C1-28** | TRAV8-3 | TRAJ28 | CAVGAPSGAGSYQLTF | TRBV4-1 | TRBJ2-7 | CASSPTSGIYEQYF |
| **KFJ5** | TRAV4 | TRAJ21 | CLVGEILDNFNKFYF | TRBV28 | TRBJ2-3 | CASSQRQEGDTQYF |
| **LC13** | TRAV26-2 | TRAJ52 | CILPLAGGTSYGKLTF | TRBV7-8 | TRBJ2-7 | CASSLGQAYEQYF |
| **D2H** | TRAV9-2 | TRAJ37 | CALDSGNTGKLIF | TRBV11-2 | TRBJ2-7 | CASTTGGGGYEQYF |

**Supplementary Table 2**: TCR rearrangement details for TCRs used for low-throughput Stitchr validation in Figure 2. Note that unless specified, all TCR alleles were *01 for their respective genes.

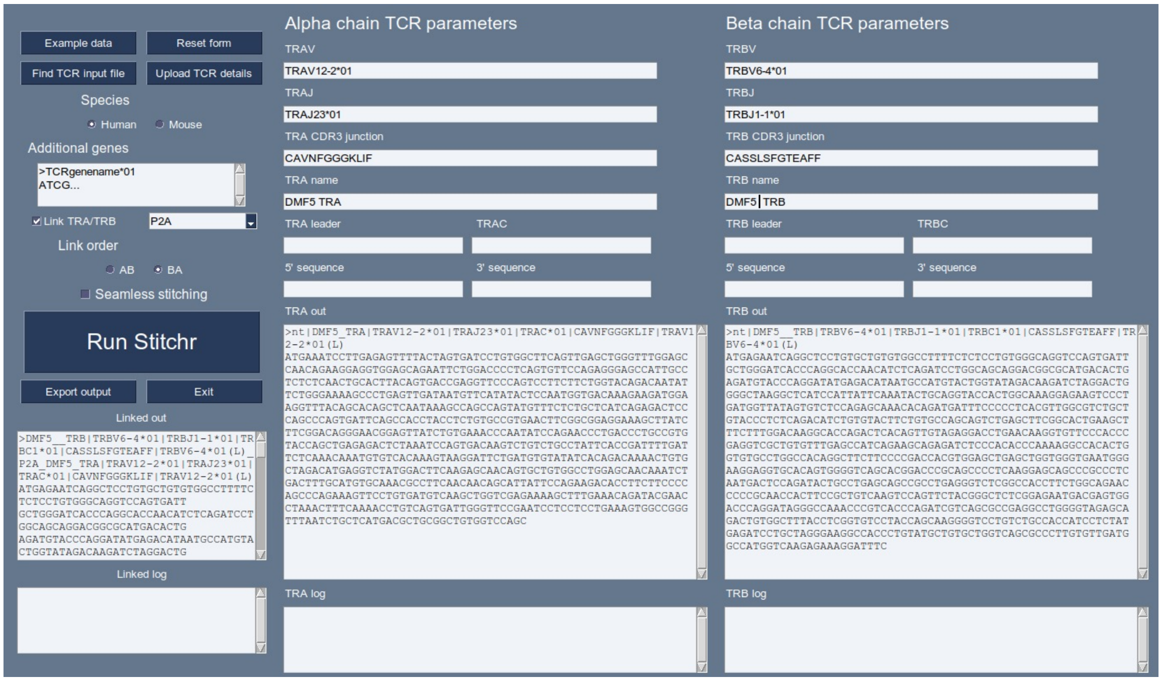


**Supplementary Figure 1.** Screenshot of the alternative graphical user interface to perform low-throughput Stitchr functions.

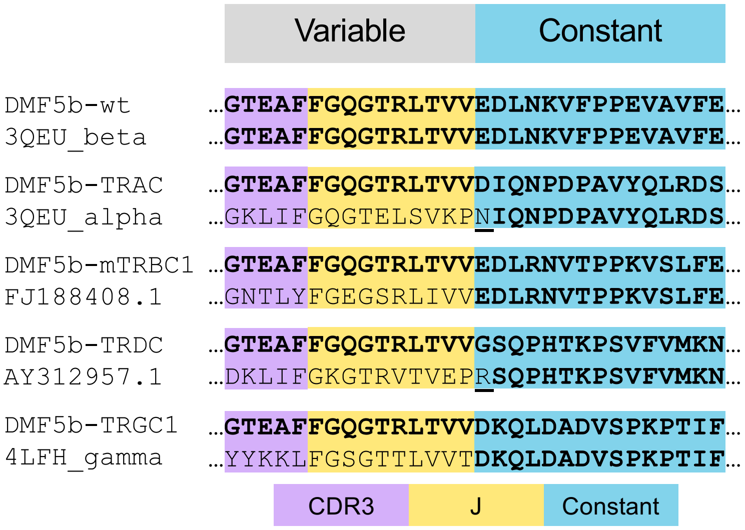


**Supplementary Figure 2. Demonstration of Stitchr's utility in TCR engineering, through constant region domain swapping.** The anti-MART1 TCR DMF5 beta chain (DMF5b) was stitched to a variety of constant regions, and then aligned against various rearranged TCRs that incorporate the same constant region. The variable:constant domain interface is shown. From top to bottom, the stitched TCR/reference pairs are: DMF5b with its own TRBC1 constant region (DMF5b-wt) against its own PDB FASTA used to identify the V/J/CDR3 information (3QEU_beta); DMF5 with the alpha chain constant region (DMF5b-TRAC) aligned to the alpha chain of the DMF5 TCR (3QEU_alpha); DMF5b with a murine beta constant region (DMF5b-mTRBC1) aligned to a rearranged mouse beta chain cDNA (GenBank accession FJ188408.1); DMF5b with the delta chain constant region (DMF5b-TRDC) and a rearranged human delta chain (GenBank accession AY312957.1); DMF5 with a gamma constant region (DMF5-TRGC1) aligned to a rearranged gamma chain TCR (PDB accession 4LFH). Sequences matching the expected wild-type DMF5b sequences are in bold. Underlined residues indicate an expected amino acid mismatch: the first nucleotide of the first codon of the constant region is donated by the last nucleotide of the J gene post-splicing, thus the amino acid encoded will vary depending on the J gene used.

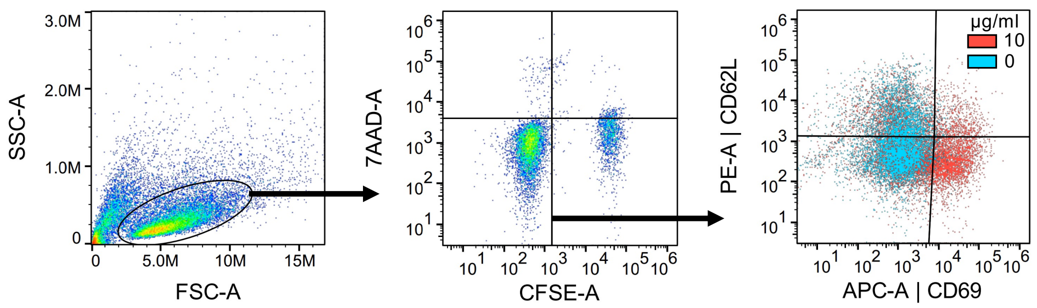


**Supplementary Figure 3. Gating strategy of TCR-transduced Jurkat activation assay**. Jurkat cells lentivirally transduced with a TCR of known reactivity were incubated overnight at a 2:1 ratio with a CFSE-labelled peptide-pulsed target cancer lines with matched or mismatched HLA-I alleles. Left to right: cells are gated away from debris, then singlets selected via FSC-A vs FSC-H (not shown), live Jurkats gated by CFSE- and 7AAD-negative. The frequency of activated Jurkats was determined by the percentage of CD69+ CD62L-negative cells.

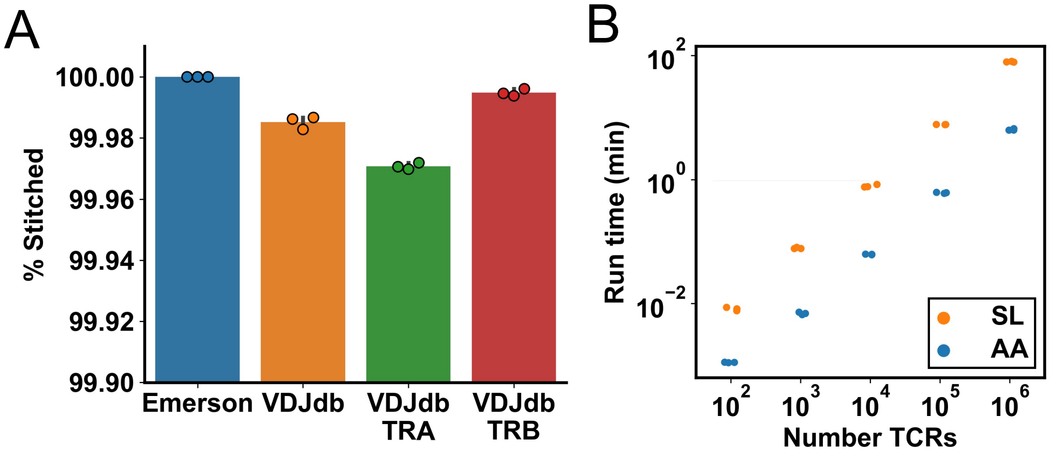


**Supplementary Figure 4. Thimble performance running on published TCR repertoire datasets**. **A,** Percentage of TCRs for which a stitched sequence was successfully produced (note cut Y axis lower bound). The Emerson data is a combination of five randomly chosen ImmunoSEQ beta chain runs, while VDJdb is the entirety of the human section of that database, further then split out into alpha (TRA) or beta (TRB) rearrangements only. Repertoires were run in triplicate, having up/down-sampled to different numbers. The results shown represent the percentage of stitched results from 1e6 TCRs. **B,** Comparison of run times for the Emerson data, up/down-sampled to different numbers of TCRs (X axis) when providing the CDR3 junction either as the junction-only amino acid sequence (AA) or the entire ~90 nucleotide sequence of the read (as listed in the 'rearrangement' column of the data) with the seamless stitching mode (SL) enabled.

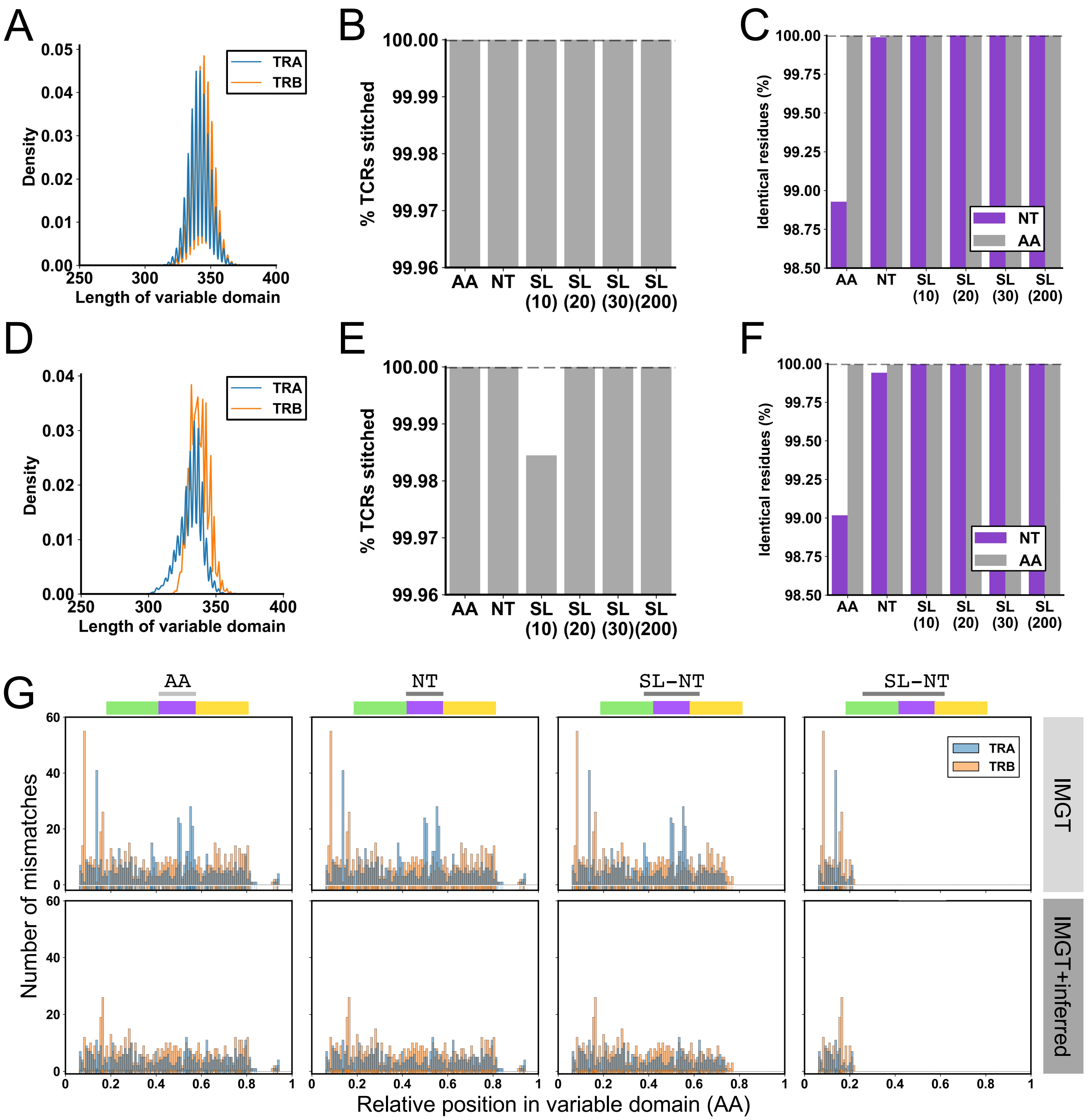


**Supplementary Figure 5. Performance of Stitchr applied to high-throughput datasets**. **A,** Density plot of variable domains (start of V-REGION to end of J-REGION) of TCRs produced by immuneSIM. **B,** Percentage of immuneSIM TCRs that successfully produced a stitched sequence. **C,** Percentage of residues (nucleotide in purple, amino acid in gray) that are identical between the input immuneSIM and output Stitchr/Thimble sequences. **D-F,** As in **A-C**, but for empirically sequenced TCRseq datasets from prior publications. **G,** Distribution of amino acid mismatches between translated sequences of TCRseq data when processed using just the typical IMGT reference database (top) versus IMGT supplemented with potential novel alleles inferred from the donors in the cohort (bottom), comparing different junction inputs (left to right: AA; NT; seamless 20-20; seamless 200-30).

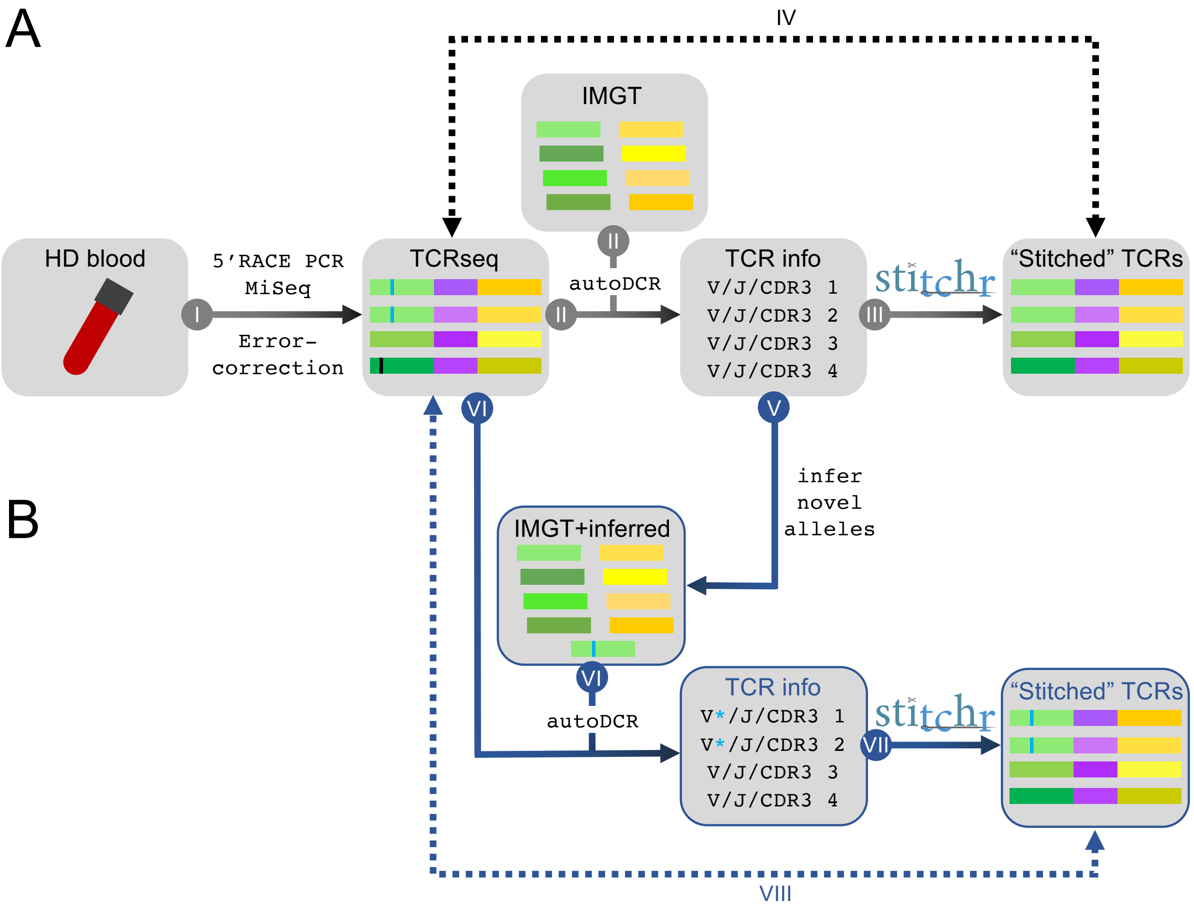


**Supplementary Figure 6. Overview of Stitchr validation on real-world high-throughput TCRseq data**. **A,** TCRs were sequenced from healthy donor peripheral blood RNA using a UMI-based 5’RACE protocol, sequenced on an Illumina MiSeq, and overlapping paired-ends were merged and error-corrected (I, see methods). Initially, TCRs in long error-corrected reads were annotated using autoDCR supplied with the IMGT reference database of V/J genes (II). TCR annotations produced were input to Stitchr (III, via Thimble), and sequences produced were compared to the sequences of the original reads used (IV). **B,** Alternatively, TCR information called by autoDCR was used to infer potential novel TCR V gene alleles which were then added to the IMGT reference (V), which were then again used to annotate the original corrected reads with autoDCR (VI). TCRs annotations produced with the IMGT+inferred reference (VII) were then compared to the original TCR sequences (VIII). Note that this process now accurately introduces SNPs contained in the novel TCR alleles (*), but will fail to replicate other mismatches between the read and the reference (e.g. sequencing errors) if they fall outside the range of a provided junction sequence.
